# The Cellular Immunity Agent Based Model (CIABM): Replicating the cellular immune response to viral respiratory infection

**DOI:** 10.1101/663930

**Authors:** Andrew Becker, Gary An, Chase Cockrell

## Abstract

Viral respiratory infections, such as influenza, result in over 1 million deaths worldwide each year. To date, there are few therapeutic interventions able to affect the course of the disease once acquired, a deficit with stark consequences that were readily evident in the current COVID-19 pandemic. We present the **C**ellular **I**mmune **A**gent **B**ased **M**odel (CIABM) as a flexible framework for modeling acute viral infection and cellular immune memory development. The mechanism/rule-based nature of the CIABM allows for interrogation of the complex dynamics of the human immune system during various types of viral infections. The CIABM is an extension of a prior agent-based model of the innate immune response, incorporating additional cellular types and mediators involved in the response to viral infection. The CIABM simulates the dynamics of viral respiratory infection in terms of epithelial invasion, immune cellular population changes and cytokine measurements. Validation of the CIABM involved effectively replicating *in vivo* measurements of circulating mediator levels from a clinical cohort of influenza patients. The general purpose nature of the CIABM allows for both the representation of various types of known viral infections and facilitates the exploration of hypothetical, novel viral pathogens.

## 0.1 Introduction

Viral respiratory infections, both seasonal and novel, are one of the most significant infectious disease challenges to the modern world. Seasonal influenza infects 5-10% of the adult population and 20-30% of children annually [1], resulting in approximately 1 million deaths worldwide, most due to severe acute respiratory infection (SARI) brought on by the influenza infection and/or secondary infections. Understanding the dynamics of viral respiratory infections is increasingly more important in light of the SARS-CoV-2 pandemic, which has resulted in 629,000 deaths worldwide as of July 24, 2020 [2] while significantly disrupting of nearly every aspect of public life. A major component of the challenge presented by SARI and viral pneumonia is the fact that there is a relative paucity of effective anti-viral agents; post-infection therapy is primarily limited to supportive care, such as supplemental oxygen and ventilation [3], as the infection runs its course. In addition to the persistent difficulty of developing anti-viral agents, the fact that disease severity can be attributed the systemic response to infection presents another challenging aspect to the search for effective treatments [4]. Characterizing the detailed underlying immune dynamics of these viral infections could facilitate the development of new therapies that modulate these dynamics in order to positively affect the course of disease. While there is promising data to suggest that immune manipulation in COVID-19, be it through steroids [5] or anti-mediator therapy [6], may be effective, there are also contradictory findings that other bioplausible anti-cytokine therapies may not be effective [7]. This pattern of promise and disappointment has marked the historical failure of attempts to modulate the immune system during an acute infection, where nearly 50 years of research have provided no currently approved pharmacological agents able to affect the underlying biology of disordered systemic inflammation (i.e. sepsis) [8]. We have suggested that the heterogeneity of clinical populations, and the sepsis population in particular, manifesting both among different individuals and within an individual’s course of disease [9], points to the need to develop precise, multi-modal therapeutic regimens personalized to individual disease trajectories with the aid of mathematical/computational modeling. In prior work we have defined the scope of interventions required to affect effective control of sepsis using a combination of mechanism-based multi-scale models (specifically agent-based models or ABMs) with evolutionary computing (genetic algorithms) [10] and model-based artificial intelligence/deep reinforcement learning [11]. This work suggests that the complex and heterogeneous nature of the clinical immune response is an example of an perspective that notes the limits of current biomedical experimental approaches for adequately exploring underlying functional relationships in human systems [12, 13]; dynamic mathematical and computational model can provide an adjunct to traditional experimental and data analytical methods to increase this understanding. In terms of viral respiratory infections, mathematical/computational modeling has been successfully used to examine multiple aspects of influenza: replicating flu dynamics by predicting viral load, response to antivirals, and immune dynamics, among other insights [14–16]. These models, which primarily use deterministic differential equations, are well suited for characterizing system-level dynamics and showing how the changes in basic system characteristics, such as virulence, viral load, and host resistance, can affect disease outcome. However, differential equation models are limited in their ability to represent the component-level stochasticity, behavioral heterogeneity and spatial effects that generate the variations in individual disease trajectories. With the development of technologies able to characterize cellular behavior at a much more detailed level, down to the behavior of single cells, effectively utilizing this knowledge calls for modeling approaches able to represent such granular information.

Agent-based models (ABMs) are discrete event, rule-based, spatially explicit computational models that have been used for decades to explore and quantify aspects of dynamic and complex systems [10, 17]. ABMs treat systems as aggregates of multiple components (“agents”) where system level behavior arises from their interactions. The ready mapping of computational agents to biological entities (such as cells) and the ability of an ABM to represent physical processes like diffusion or force applications facilitate the use of ABMs to model various biological systems, such as the immune response. ABMs have been used to investigate acutely disordered systemic inflammation, otherwise known as sepsis, [9, 10]. Sepsis is known to involve a hyper-inflammatory response, or ‘cytokine storm’ [18], on top of systematic viral or bacterial infection, and as such prior ABMs of sepsis may be relevant in the search for effective treatments for SARS-CoV-2 infection, which is also theorized to have cytokine storm as a leading cause of mortality [19]. While our previous models investigating sepsis primarily represented bacterial infections and the innate immune system, the response to viral infections typically requires the adaptive immune response in order to completely clear an infection [20]. As such we need to extend these original innate immune ABMs to include aspects of adaptive immunity.

In the current work, the simulation takes place on a stationary/fixed portion of pulmonary epithelial tissue; agents (immune cells, blood cells) traverse this grid and interact with it and each other through the secretion of cytokines or propagation of a pathogenic infection. We augment previous models of innate immunity through the addition of CD8+ T-cells, immune memory, and myeloid dendritic cells. Additionally, we augment those models by compartmentalizing epithelial cells, endothelial cells, primary lymphatic tissue (PLT), and secondary lymphatic tissue (SLT). This allows us to then calibrate the model to a specific type of infection for model validation. Herein we use cytokine levels drawn from McClain et al. [21] as a baseline data set, along with qualitative analysis of cell population and cytokine dynamics during and after an infection.

## 0.2 Materials and Methods

The current model, termed the Cellular Immunity Agent Based Model (CIABM), is an extension of the previously validated Innate Immune Response ABM (IIRABM) from Cockrell and An [9]. The CIABM was implemented in C++; the code for the CIABM is available at https://github.com/An-Cockrell/CIABM_release. In order for the CIABM to represent viral infections, the following new agents were added: myeloid dendritic cells (mDCs), naive CD8+ T-cells (Naive T), cytotoxic CD8+ T cells (Cyto T), T Effector Memory cells (TEM), T Central Memory cells (TCM), and **T**erminally-differentiated **E**ffector **M**emory T cells **R**e-expressing CD45**A** (TEMRA).

### 0.2.1 Topology and System-Level Features of the CIABM

The CIABM is a two-dimensional abstract representation of human endothelial-blood-tissue interface. This abstraction is designed to model these interfaces during an infection and does so by representing these interfaces as the unwrapped internal vascular surface of a 2D projection of the capillary vascular network. The closed surfaces can be represented as a torus, and these two-dimensional surfaces define the interaction space simulated by the model. Organ interfaces are represented via several compartments which are separated by either fully distinct 2D surfaces or by tags within the model.

The CIABM is divided into two interfacing tori. The first torus represents the surface of the circulatory system with 3 segments in series each representing a particular functional compartment: the vascular capillary network of endothelial cells, the primary lymphatic bed, and the secondary lymphatic bed (see Figure 1). This torus interfaces with a second torus, representing tissue or epithelial cells, that maps onto the capillary/endothelial segment of the circulatory torus; circulating immune cells can migrate “through” the endothelial surface and into the tissue/epithelial torus. Each compartment has the same area of 101*101 tissue cells. Each location on this surface contains a static agent which preforms the function of the segment (i.e. endothelial, primary and secondary lymphatic, and epithelial cells). Agents which represent various immune cells can travel across these surfaces. The edges connect forming a toroidal geometry for the two surfaces, one 101*101 cell-widths and one of 303*101 cell-widths, as shown in Figure 1. It is important to note that the boundaries between tissues on the larger torus are not freely crossed, but only crossed when specific signals are present (See Table 1 in Supplemental Material). While not pictured the smaller torus is effectively ‘sleeved’ in the Z-axis around the endothelial tissue as travel occurs between those surfaces.

**Figure 1:**
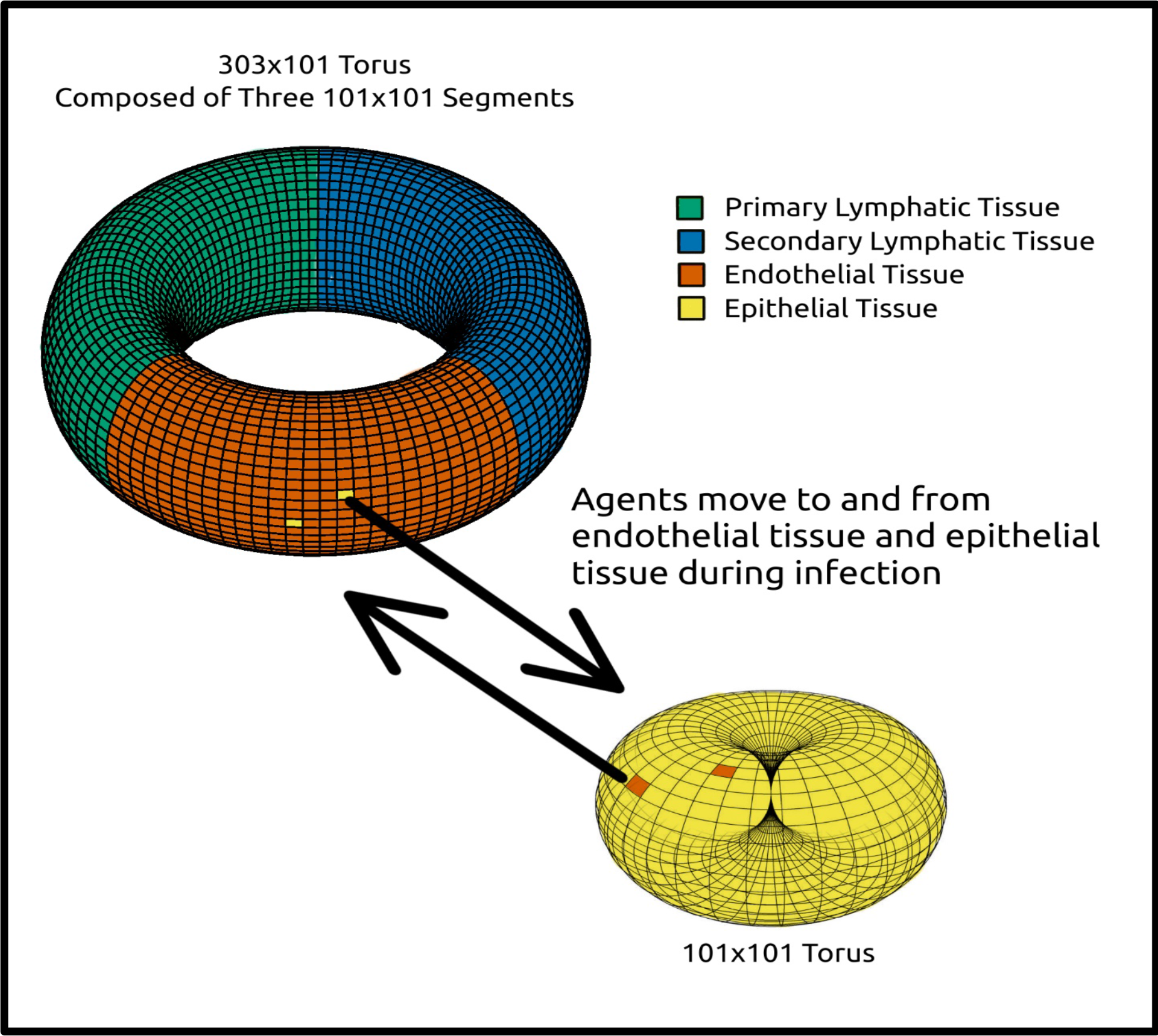
Diagram of tissue surfaces folded into tori and transfer of agents between those tori

Cellular agents can travel either around the 303*101 section of tissues or the 101*101 section of epithelium. Each of these two sections have periodic boundaries which form the tori. Some cellular agents can also travel between these major compartments when the endothelium adjacent to the epithelium becomes permeable to cells and cytokines. Cytokines can also diffuse between these surfaces. This behavior is stimulated by IL-1 or IFNα being present in the extracellular fluid between these two tissues. These cytokines are generated in direct response to tissue damage, viral infection within the epithelium, or exposure to extracellular virions, each of which can then initiate a cascade of reactions simulating the innate and adaptive immune response.

Cytokines, viral particles, and toxins are represented as positive-valued variables within every tissue cell. Each stationary tissue cell holds information regarding the concentration or count of each of these variables at its Euclidean coordinates. Each cytokine has a specified rate of diffusion and degradation (‘evaporation’), representing the array of non-cellular effects which contribute to a protein half-life, ultimately representing the true rate of elimination of each compound within the human body [22]. There is additionally a constant rate of diffusion of each cytokine across the semipermeable endothelium during inflammation. This method of storing information allows for agents moving across the simulation to modify and access these values at any point without storing any information about them. This results in a nearly memoryless model, where the current state of the model and each step dictates the actions of each cell. The exception to this ‘memorylessness’ is the development of immune memory, which stores information about the antigen of prior infections explicitly in the adaptive immune cells which proliferated in response to that infection.

Viral particles in the extracellular fluid are also represented as environmental variables (and not as individual agents), with specifically defined diffusion/spreading rates and a rate of ‘evaporation’, representing the removal, destruction, or inactivation of viral particles in blood and extracellular fluid. These rates are modulated by cytokine concentrations within the system at each cell location. As the simulated infection in this work is influenza, viral particles can infect and replicate only one cell type in our model, the epithelial cell; immune cells do not become infected. Viral particles in the extracellular fluid can enter epithelial cells and replicate, but do not replicate in extracellular fluid. The rate of entry is probabilistic and calculated for each epithelial tissue cell at each time step. These particles only increase in number within these cells (representing intracellular viral replication) and are only released when a critical mass of particles is reached, leading to a simulated viral burst based on actual (*in* vivo) viral burst counts in similar viral infections, [23].

Epithelial cell death can occur under three conditions: 1) when the virion count within the cell reaches critical and a viral burst occurs, 2) when a cell has been infected for a long time without reaching critical mass and autolyses, or 3) when cytotoxic compounds are present in sufficient concentration to kill the cell. There is interplay between cytotoxic compounds, immune cell interaction, and viral infection in determining the number of time steps before a cell autolyses. When a critical mass of cells has been destroyed the system is dead and the simulation stops: this threshold is arbitrarily chosen as 50% of the cells. Note that epithelial cells will also heal over time if three conditions are met: 1) it is not infected, 2) there is a lack of inflammatory cytokines, and 3) adjacent cells are healthy. For each cell it takes approximately 5 days without additional damage for the cell to heal [24].

Rate of cytokine production, viral replication, viral destruction, cell damage, cell activation, and other activities are controlled by a rule matrix. This rule matrix, **M**, follows one of two formulas to calculate the new value of the variable being modified. The first of these is a *production rule*, which constrains the result of the operation by the cell to always be greater than or equal to zero. For a production rule the value of every concentration variable within the system is loaded into a vector, **C**. We then select the appropriate row from **M** which corresponds to the rule being accessed, which we call **R**. Then by multiplying **R** x **C** we calculate the change in the value in question, *A*. For the production rule we ensure this value is 0 or greater by assigning the value 0 in the case where *A* < 0. This indicates that signals that inhibit the synthesis of that specific protein dominate the interaction over those that promote the protein synthesis. The second formula, a *combination rule*, does not apply this last step of reassigning sub-zero values to zero, and allows for negative output.

These base dynamics are augmented by each cell type within the system. General cell dynamics, and then specific cell types and their rules are outlined in the rest of this section.

### 0.2.2 Generic Cell Dynamics

The CIABM contains three broad categories of cells:

1. Static cells, which occupy grid spaces in the ABM world and do not move; in the current model these represent endothelial cells, which line blood vessels, and respiratory epithelial cells.
2. Progenitor cells that reside in specific locations within the model and produce additional cells of specific types. These progenitor cells represent abstractions of the hematopoietic lineages that produce immune cells and include PMN progenitors, TH0, TH1, and TH2 progenitors. These progenitors are the only source for new immune cells, with the exception of naïve CD8 T Cells, which can produce copies of themselves upon recurrent stimulation by the same antigen.
3. Active cells, which represent all the mature immune cells. Active cells travel within the model from region to region, produce and react to cytokines, and perform the effector functions of the immune/inflammatory response.

Active cells have the most complex rules. These rules involve movement and orientation of the cell within the system, transfer across borders, location detection, and gradient descent. Each of these cell types contains information regarding the location of the cell in the form of an x, y coordinate pair, the tissue type the cell is residing in, and orientation of the cell within the grid. Orientation within the simulated environment uses an 8-point scale, each cardinal and ordinal direction on a compass, to label the direction the cell is going to head upon the next movement within the system. Cellular movement is determined by mediator gradients, which has been observed as increased chemotaxis at low concentrations of compounds, and reduced chemotaxis at high concentrations [25].

### 0.2.3 Individual Cell Types and Functionality within the CIABM

The CIABM has a total of ten active cell types, seven progenitor cell types, and two tissue cell types. Each of these cell types has unique rules and functions. Figure 2 shows a wireframe diagram of a simplified version of these rules; if all interactions between cell types and all progenitor cells were to be included the resulting figure would be very difficult to parse without adding informative data. In this diagram we have excluded several compounds which are not currently used in control loops and most progenitor cells. Additionally, only strong feedback from compounds is included as red lines in the model. The feedback included in Figure 2 drives the overarching patterns of feedback within the model.

**Figure 2:**
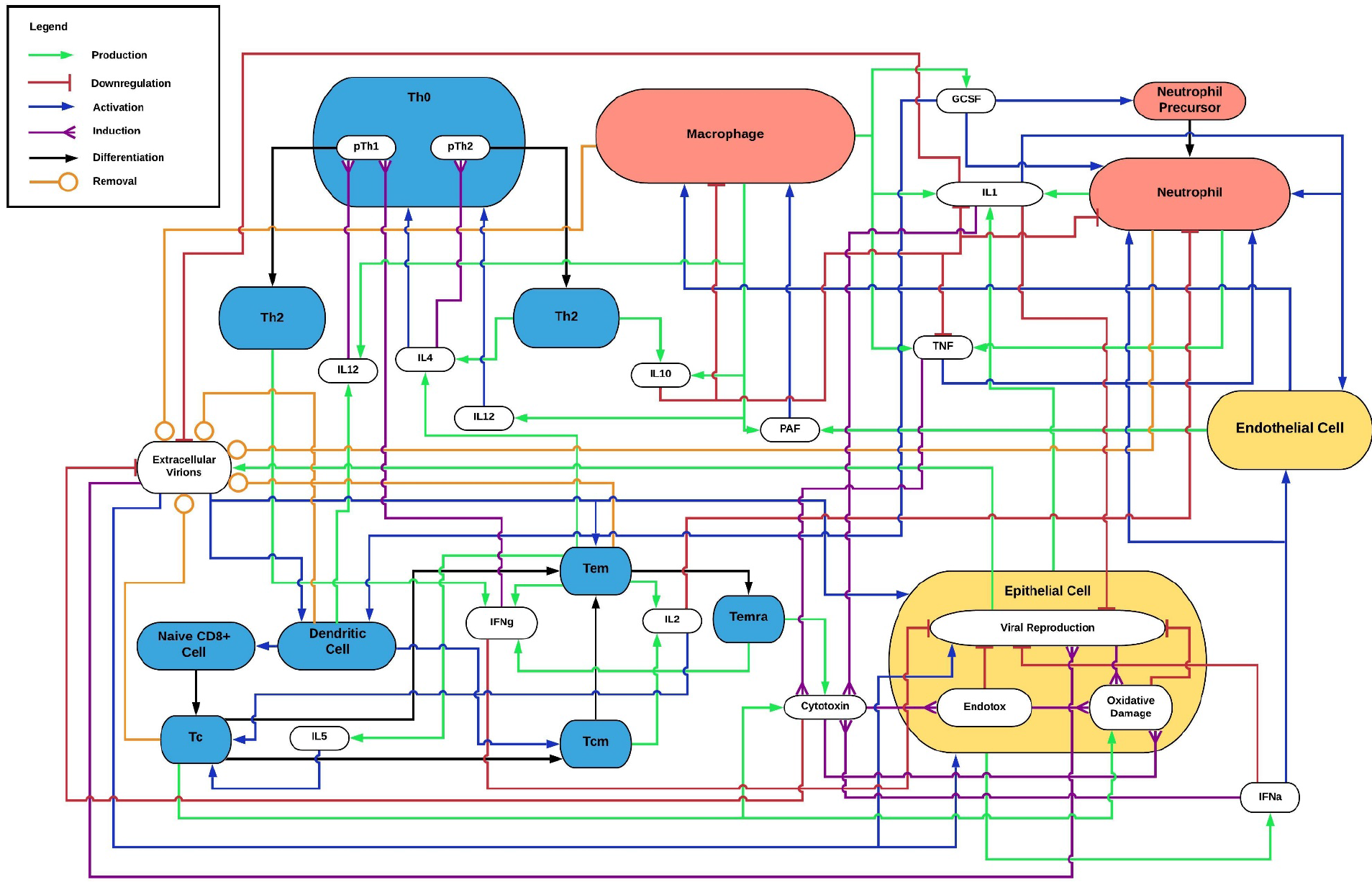
Simplified diagram of interactions between cell types, compounds, and virions

Even in the reduced set of interactions depicted in Figure 2 there are several multi-level control loops evident, where a single stimulus, such as the presence of viral particles, or the introduction of a particular compound, can set off a self-extinguishing chain reaction which eliminates that compound or insult. There are only two outcomes from any perturbation of the system: either the system finds an equilibrium or there is an overwhelming amount of damage to epithelial cells. Perpetual active infections are rare in our model and tend to collapse into eliminating the infection completely or system death within several simulated weeks.

#### Innate Immune Cells

There are five cells that model the dynamics of the innate immune system: the two types of tissue cells, myeloid dendritic cells, monocytes and polymorphic nuclear cells (PMNs).

Epithelial tissue acts as the first defense in the innate immune system of the lungs. If the virus cannot get a foothold in the epithelial tissue, then infection does not occur. This defense is represented as a probabilistic entry chance for viral particles at each tissue location during each time step, and the fact that viral particles get eliminated from extracellular fluid at some rate each turn. If the initial viral load is extremely low than it is possible no infection of the cells occurs at all, and in turn no immune memory is developed. Once epithelial cells are infected they begin to produce interleukin-1 (IL-1), tumor necrosis factor alpha (TNFα), and interferon alpha (IFNα), which is produced by lung epithelial cells during influenza infection [26]. The rate of production is determined from current concentrations of all cytokines, virions, and other compounds, multiplied by constant values in the rule matrix **M** as described in 0.2.

Some of the signals produced by epithelial cells act as chemo-attractants for other immune cells. TNFα promotes chemotaxis at low concentrations for polymorphonuclear leukocytes (PMNs) and reduces chemotaxis at high concentrations [25]. This results in PMNs migrating up TNFα gradients to the site of infection as described in section 0.2.2. Once at the site of infection PMNs also produce IL-1 and TNFα, which compounds this attractant effect.

IL-1 in the extracellular fluid can trigger endothelial cells to become permeable to immune cells, various compounds, and infectious agents, and additionally to produce another set of cytokines. In our model once activated endothelial cells produce platelet activating factor (PAF) and interleukin-8 (IL-8) [27, 28]. PAF functions as a chemoattractant for monocytes [29].

Monocytes then perform the same gradient descent in the direction of the activated endothelial cells, and then migrate into the site of infection. Once activated in the site of infection, monocytes make a large suite of compounds which are outlined in Table 1 in supplemental material. Among these is granulocyte colony-stimulating factor (GCSF), which functions as a chemoattractant for mDC’s and stimulates the recruitment of PMN’s. Myeloid dendritic cells act as the bridge between the innate immune system and the adaptive immune system and are an essential component of this connection [30]. They migrate according to the gradient of a suite of inflammatory cytokines, uptake infectious material, and then travel out of the site of infection and to the lymph node. Once in the lymph node mDC’s present this antigenic material to naïve CD8+ T Cells. In the presence of the correct signals, and with enough contact, this results in the beginning of the adaptive immune response.

Both monocytes and PMN’s have progenitor cells within the model which maintain their populations during infection. These reside in PLT and are activated by GCSF diffusing throughout the system and produce more monocytes and PMNs. These newly produced cells migrate into the blood, and if the correct signals are still present, will migrate into the site of infection [31]. mDC’s have progenitor cells but they produce mDC’s at a constant rate regardless of infection, and the mDC’s age and die regardless of infection. This reflects the data showing that mDC populations do not change during infection and their limited lifespan [32, 33].

#### Adaptive Immune System

The adaptive immune system in our model is the suite of cells associated with developing immune memory and modulating the immune response as the response and infection progress. This includes two groups of cells, T helper cells, which modulate the response, and CD8+ cells, which eliminate any remaining infection caused by specific pathogens and maintain immune memory to those pathogens. This separation of tasks allows for fine tuning of the model and the effects of each cell. Each of these cells also produces specific cytokines and interacts with other cells in specific ways.

T helper cells, also known as CD4+ T cells, help to modulate and direct the immune response. In our model we include progenitor cells which reside in the PLT and stochastically produce our three types of helper cells: T-helper 0 (Th0), T-helper 1 (Th1), and T-helper 2 (Th2) cells. This results in constant replacement of these cells and a consistent total population in a similar way to mDCs. As seen in Figure 2 Th1 and Th2 cells modulate innate and adaptive immune cell populations for cellular succession to occur [34]. Th0 cells develop into Th1 or Th2 cells differentially depending on the concentration of various cytokines, which change over the course of infection. The population switches to become majority Th2 as the course of infection progresses, and these cells downregulate innate immune cells through cytokine production while upregulating themselves by ensuring a larger proportion of Th0 cells become Th2 cells. Th1 cells also self activate and help to modulate the initial adaptive immune response through production of interferons [35].

The CD8+ T cell response in the most complex cell succession within this model. In order to initiate the CD8+ T cell response in a naïve immune system, one which has not experienced a particular antigen, an mDC must travel from the site of infection with an antigen from that infection and find a naïve CD8+ T cell with a receptor for that specific antigen. In our model we populate the SLT with naïve CD8+ T cells that have a random number between 0 and 9 that represents the antigen it can bind to. Each infection also generates a random number between 0 and 9, and when an mDC collects antigenic material, our model represents that as holding the number associated with that pathogen. Once the mDC encounters a naïve CD8+ cell with the proper cognate antigen, they bind together and begin the process of proliferation [36].

During proliferation naïve CD8+ cells produce Cyto T, Tem, and Tcm cells. Naïve CD8+ cells also produce copies of themselves. All cells have the same cognate antigen as the parent cell, and the majority are Cyto T cells. Cytotoxic CD8+ T cells are the primary effector cells of the adaptive immune system during a viral infection. They produce granzyme and other compounds which attack virally infected immune cells and destroy viral particles [37]. Cyto T cells have a limited lifespan, and at the end of that lifespan they either die or proliferate into either a Tem or Tcm cell. This is dependent upon the concentration of various cytokines [38]. Cytokines produced later in infection bias the cell towards production of Tcm cells in our model.

Tem cells and Tcm cells are long-lived memory cells that reside in the tissue and SLT respectively. These cells can be reactivated quickly upon an encounter with their cognate antigen. Tem cells function as a population of long-term effector cells which reside in the tissue and augment the initial innate immune response, whereas Tcm cells reside in the SLT and provide a quicker source of producing more T effector cells [39, 40]. Tcm cells are extremely long lived if not activated, whereas Tem eventually proceed further down the CD8+ T Cell lineage into Temra cells [41]. Tem develop into Temra cells with a fixed probability in our model. Temra cells also provide some effector function and reside within the tissue in our model [42].

These interactions are outlined in Figure 2, and you can follow the black lines from Naïve CD8+ T-cells to TEMRA cells and see the cytokines that provide feedback to the development process of these cells. Additionally, you can see how T helper cells produce some of these cytokines and modulate that development. mDC cells also are present and you can track the effect of these individual cells in Figure 2 as well

The production rules, activation rules, and proliferation rules of all cell types and pathogens have metaparameters which govern the perturbation of the system. These meta-parameters were hand tuned in order to achieve a model which accurately represents the human immune system. Meta-parameters governing pathogen replication rate and pathogen susceptibility to host factors are then tuned to provide varying courses of infection.

## 0.3.0 Results

After application of a simulated viral infection, the model executes until the system fully recovers to a state where either all epithelial cells are repaired and the infection has been cleared, or dies; as noted above we have arbitrarily defined “death” as occurring when over 50% of the epithelial cells have been killed/destroyed. The full recovery time ranges from three to five weeks depending on severity of infection. Four different levels of infection were simulated by altering specific viral properties in the meta-parameter matrix; 1) viral reproduction speed was increased, and 2) virions were rendered to more resistant to destruction by reactive oxygen species produced by immune cells. It is important to note that the rule structure internal to the model was not changed Adjusting these two factors produced simulated infections that represented: 1) mild or asymptomatic cases, 2) moderate infections, 3) severe infections, and 4) lethal infections.

In Fig. 3 we see the changes in cell populations during a mild infection. Cell damage increases with viremia, and as viremia subsides, cell damage is healed. The innate immune system, primarily PMNs, enter early and contain the infection. After several days Cytotoxic CD8+ T cells develop and migrate to tissue and remove any remaining infection. After infection subsides CD8+ immune cell populations increase and develop memory against that infection. Note that adaptive immune populations are counted in aggregate, whereas innate immune populations are only counted within the epithelial tissue, as innate immune populations are nearly static within the model while adaptive immune cells have dynamic populations in the model. Fig. 4 shows the course of selected cytokines during that same infection. In this relatively trivial infection, the systemic inflammatory cytokine response is significantly less compared to more serious infections (Fig. 5–7) and would likely present asymptomatically in the clinic. Each of these tracks is scaled for clarity such that the pattern of each cytokine and cell type is discernable within the plot. Viremia and oxygen deficit are not scaled and represent the true values within the model.

**Figure 3:**
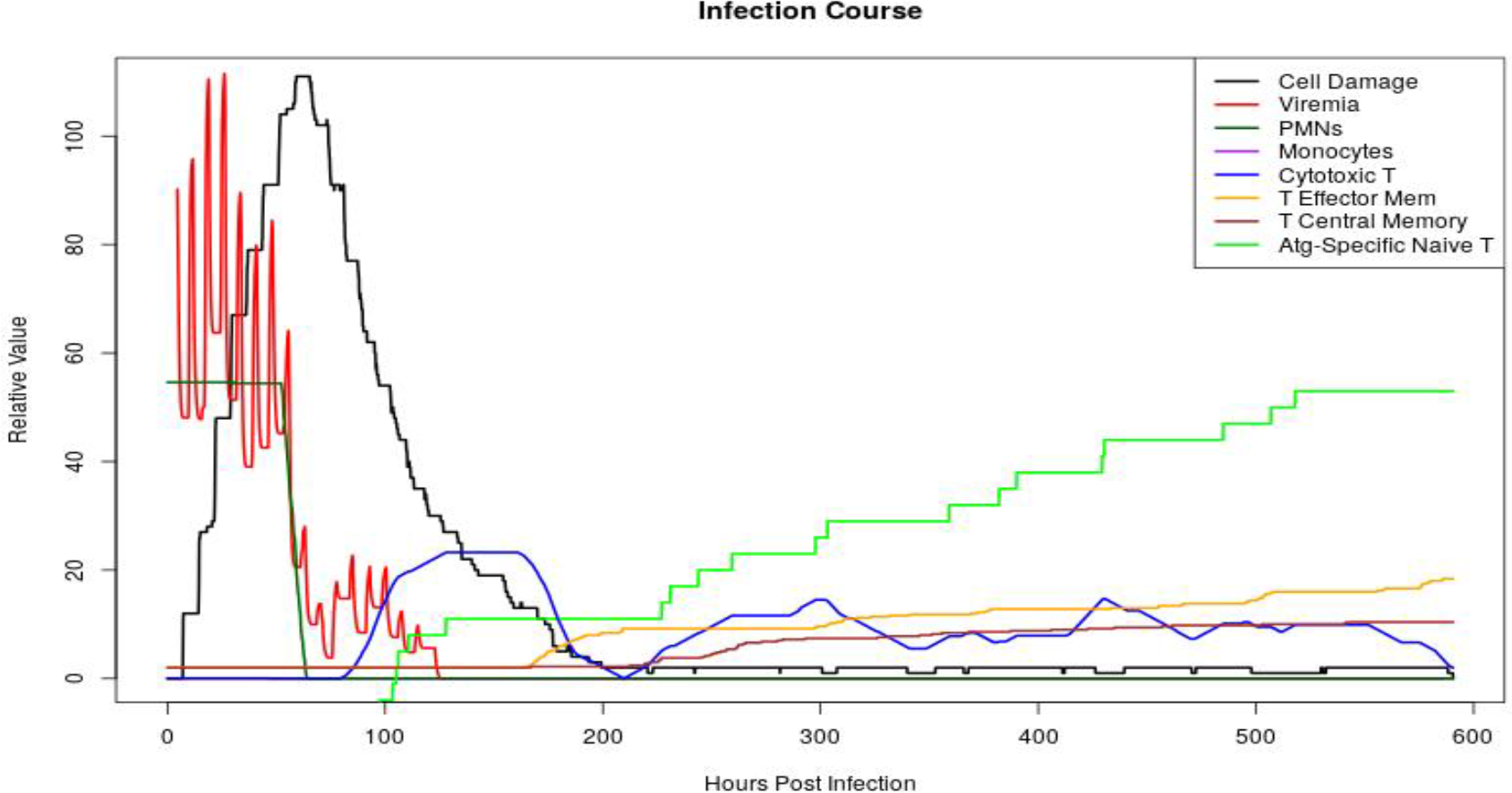
Plot of immune cell populations during the course of a mild infection

**Figure 4:**
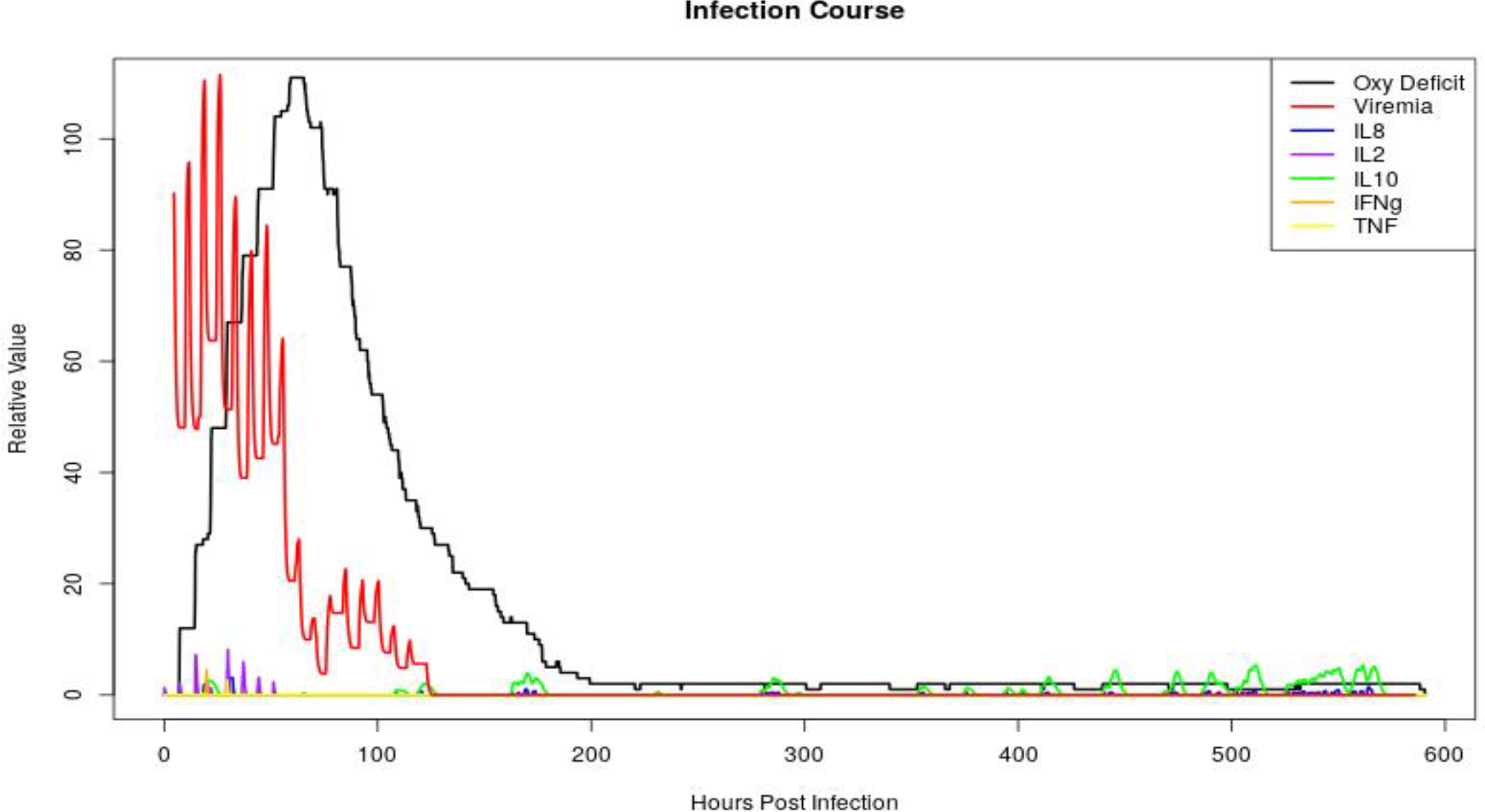
Plot of cytokine concentrations during the course of a mild infection

**Figure 5:**
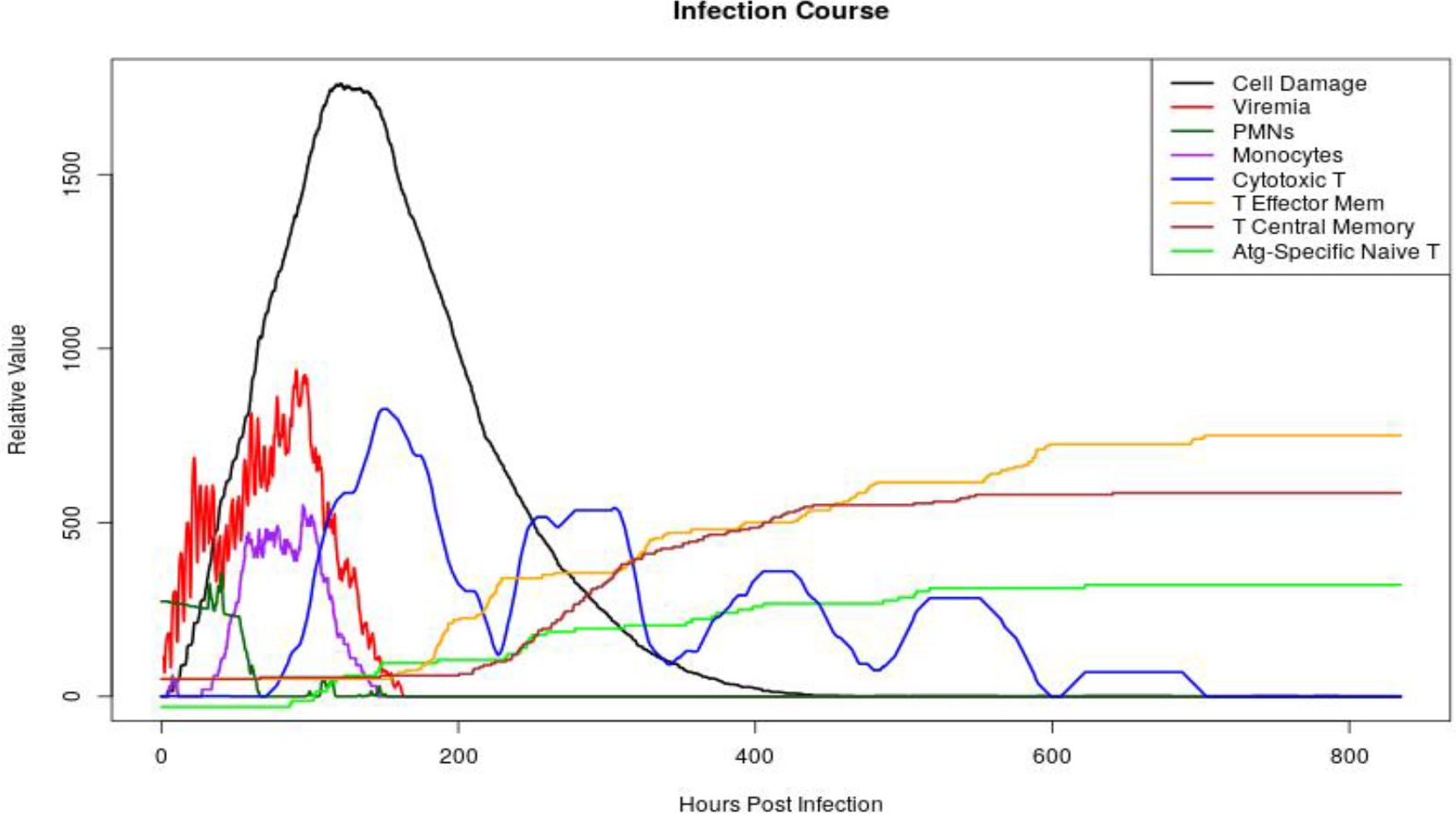
Plot of immune cell populations during the course of a moderate infection

**Figure 6:**
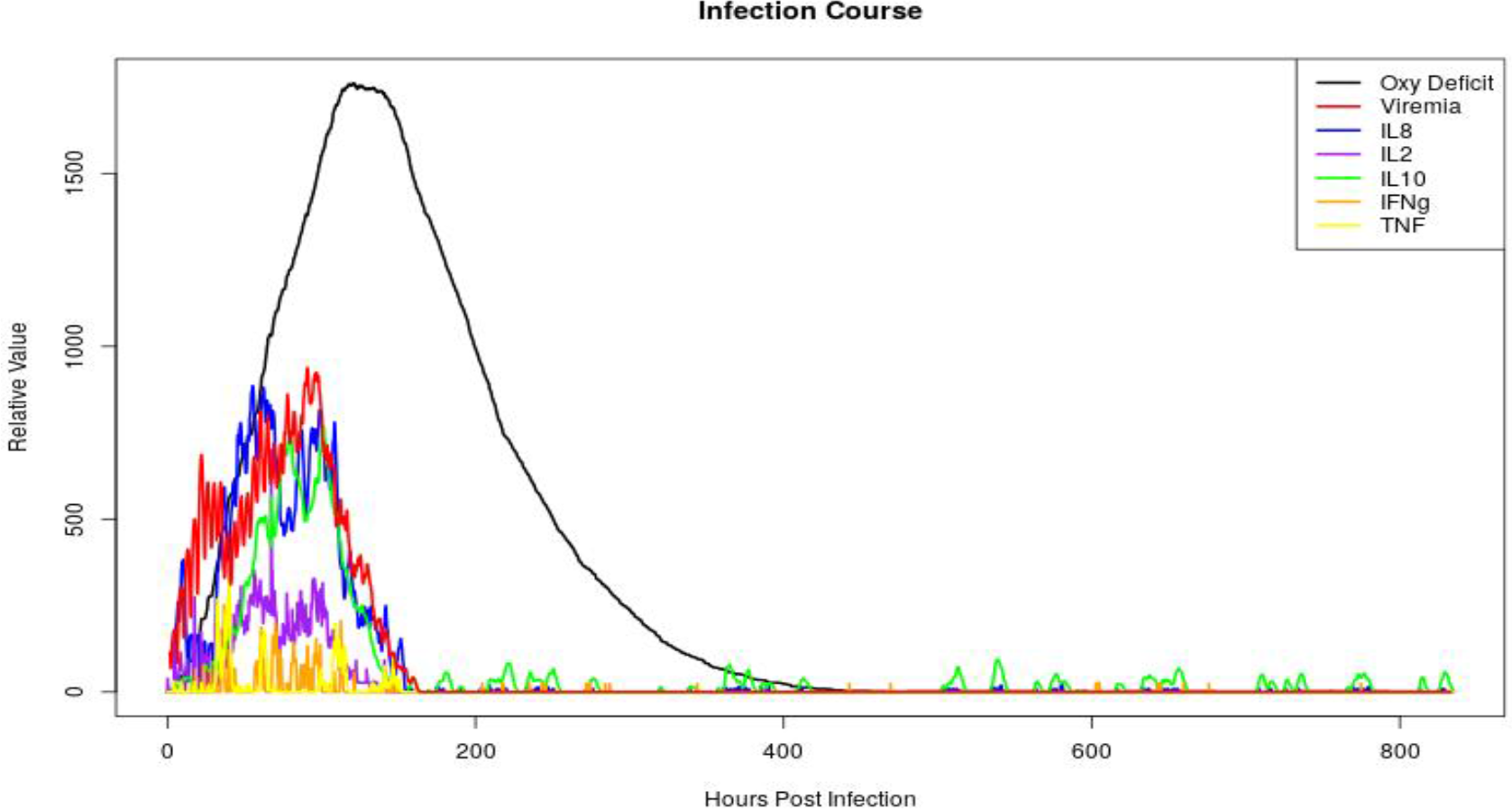
Plot of cytokine concentrations during the course of a moderate infection

**Figure 7:**
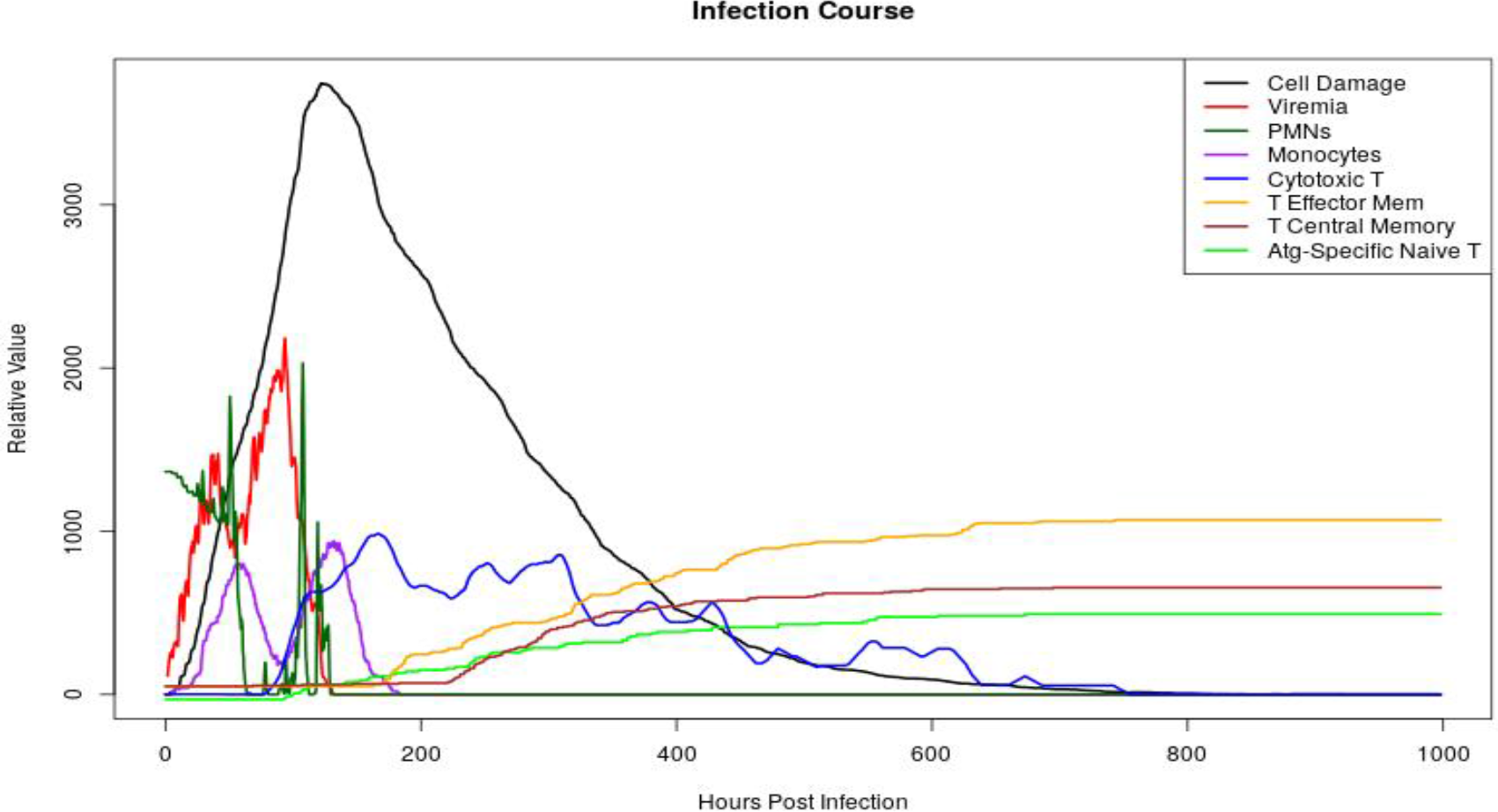
Plot of immune cell populations during the course of a severe infection

Fig. 5 and 6 present the same time-series information as in Fig. 4, however with a more serious infection: a ‘moderate infection’, which would rarely result in the death of the simulated system but would still generate a symptomatic response. In Fig. 5 we see a peak of monocytes which is not visible in the mild infection. The simulation follows the same general infection course as with the trivial infection, where the infection is contained and slowed by innate immune cells, PMNs and monocytes, and later fully eliminated by cytotoxic t-cells. Waves of cytotoxic t-cells are produced during and after infection. Post-infection there are increases in adaptive immune memory cells to retain some cell-mediated protection from this infection. In Fig. 6 cytokine signaling is more pronounced, with IFNγ and TNFα peaking early, followed by IL-2 and IL-8 which follow closely with viremia, and IL-10 peaking as viremia decreases. Small peaks of IL-10 recur as the infection resolves and the cells fully heal.

Fig. 7 and 8 present the time-series for a ‘severe infection,’ which often results in the death of the simulated system. In Fig. 7 we note that even with a robust innate immune response the ability of the infection to spread is minimally retarded and does not begin to resolve consistently until the arrival of cytotoxic t cells. There are two peaks in the viremia measurement, one before the arrival of a second wave of PMNs, and one before the arrival of cytotoxic CD8+ T cells. After the elimination of the infection immune memory population dynamics present similarly to previous infections, but to higher degree, with waves of cytotoxic t cells developing into long term central memory and effector memory cells along with continued proliferation of antigen specific naïve t cells. In Fig. 8 we see a more pronounced version of the cytokine dynamics seen in Fig. 6. There is additionally a large secondary peak for TNFα and IFNγ that coincides with the arrival of a second wave of PMNs. IL-10 is maintained at a low level as immune activity dissipates.

**Figure 8:**
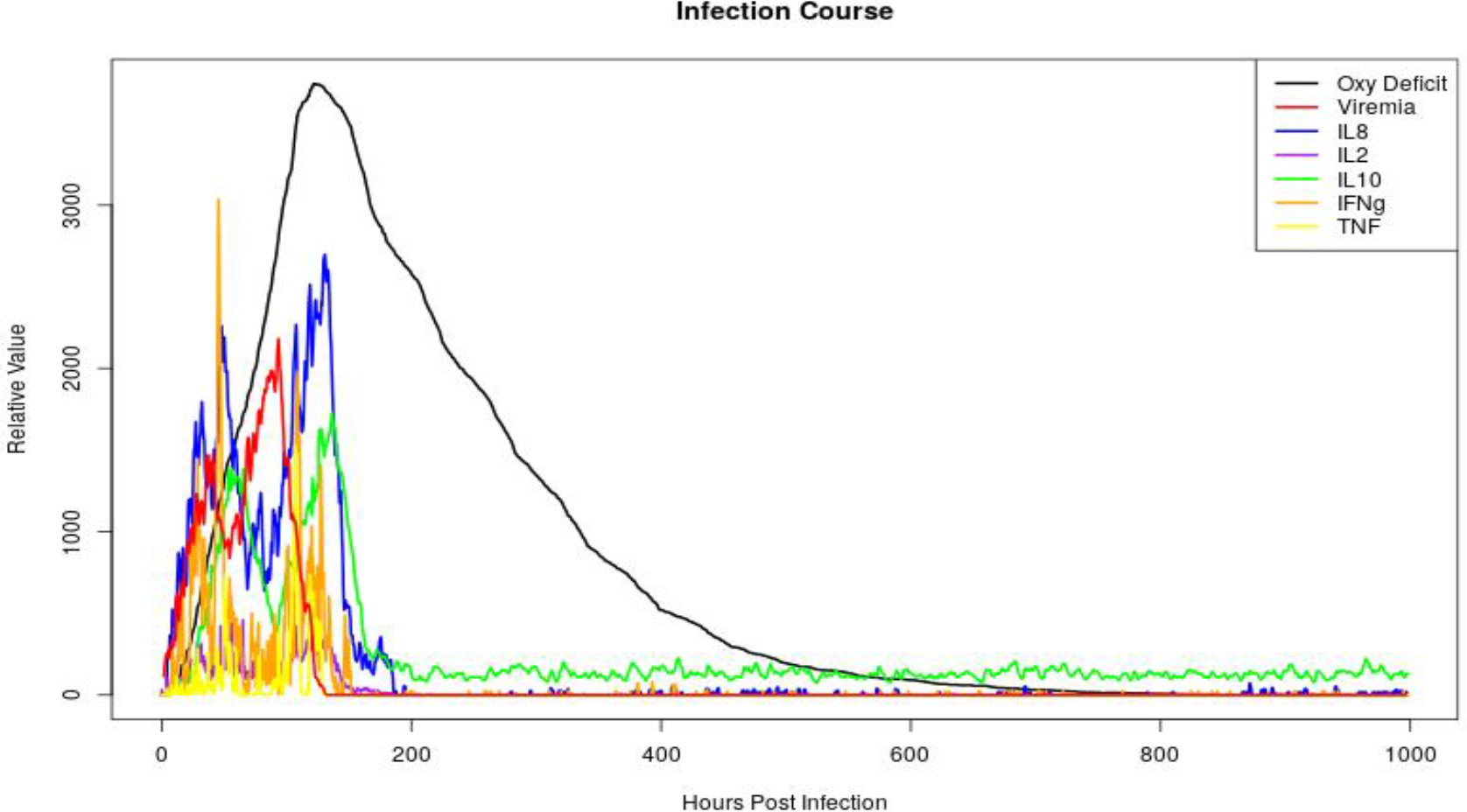
Plot of cytokine concentrations during the course of a severe infection

Fig. 9 and 10 present disease trajectories for a ‘lethal’ infection, which almost always results in the rapid death of the simulated system. In Fig. 9 we see a strong innate immune response, but the infection manages to reach the system death conditions prior to the arrival of cytotoxic T cells. Fig. 9 displays a dynamic population of monocytes and PMNs increasing with increased viral load manage to control the rise viremia, but cell damage continues to increase prior to healing, resulting in system death. In Fig. 10 we get a closer look at the pre-cytotoxic T cell dynamics of the cytokines in these figures.

**Figure 9:**
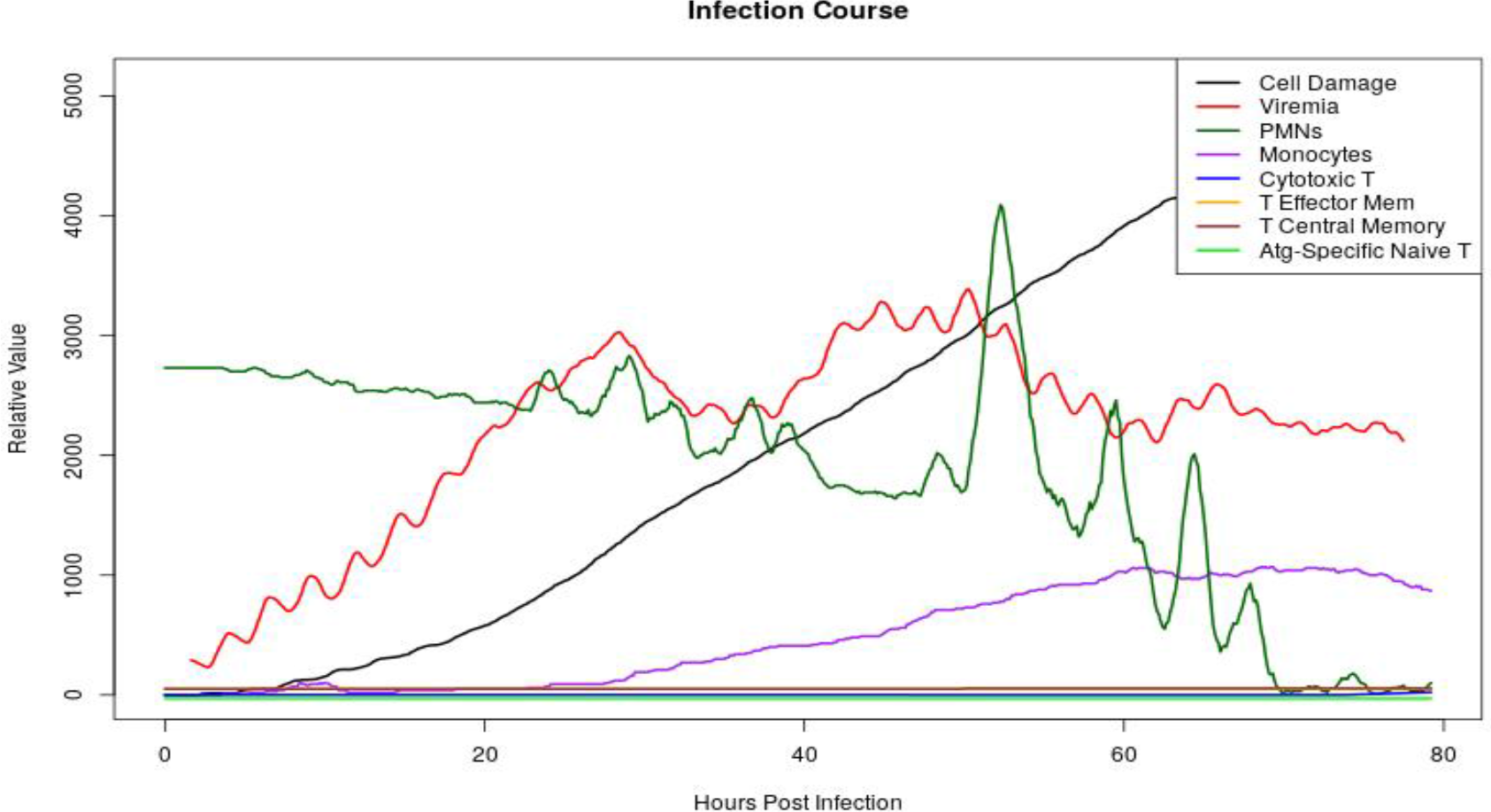
Plot of immune cell populations during the course of a lethal infection

**Figure 10:**
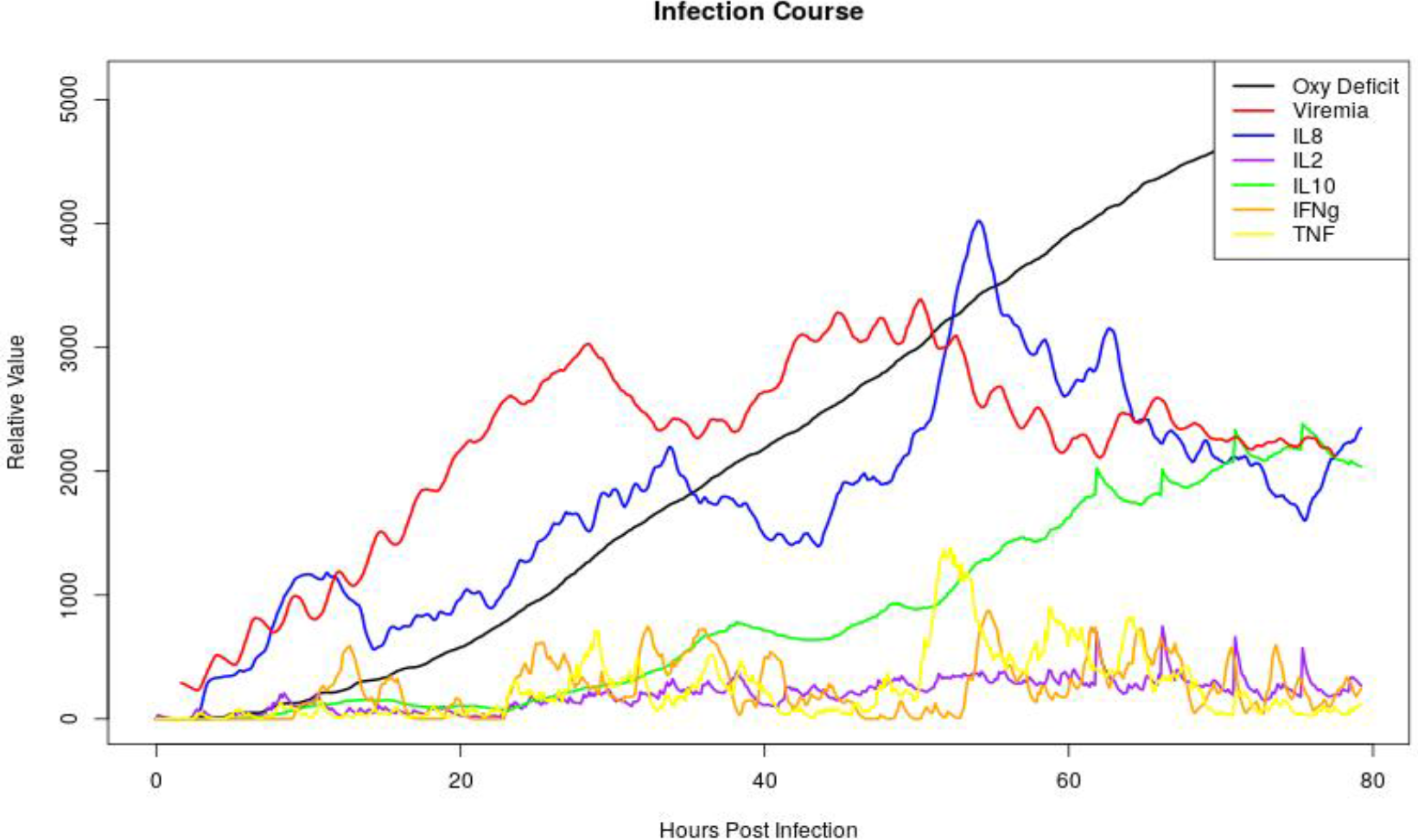
Plot of cytokine concentrations during the course of a lethal infection

Fig. 11 contains comparisons between cytokine courses in the first 1400 turns of the moderate infection and clinical measurements from McClain et al [21]. The black line is the clinical time course and the red dots are averages across 10 stochastic replicates of the CIABM using the rule matrix that produces “moderate” infections, (see Fig. 3 and 4). Error bars represent a 99.7% confidence interval, and simultaneously a 90% prediction interval, meaning we are 99.7% confident that the mean lies within that interval and 90% confident that the observed value for an addition run would fall within that interval. The clinical measurements are contained within the simulation error bars in nearly every instance. We note that information regarding the variance in the clinical data was unavailable, and thus a direct comparison is impossible.

**Figure 11:**
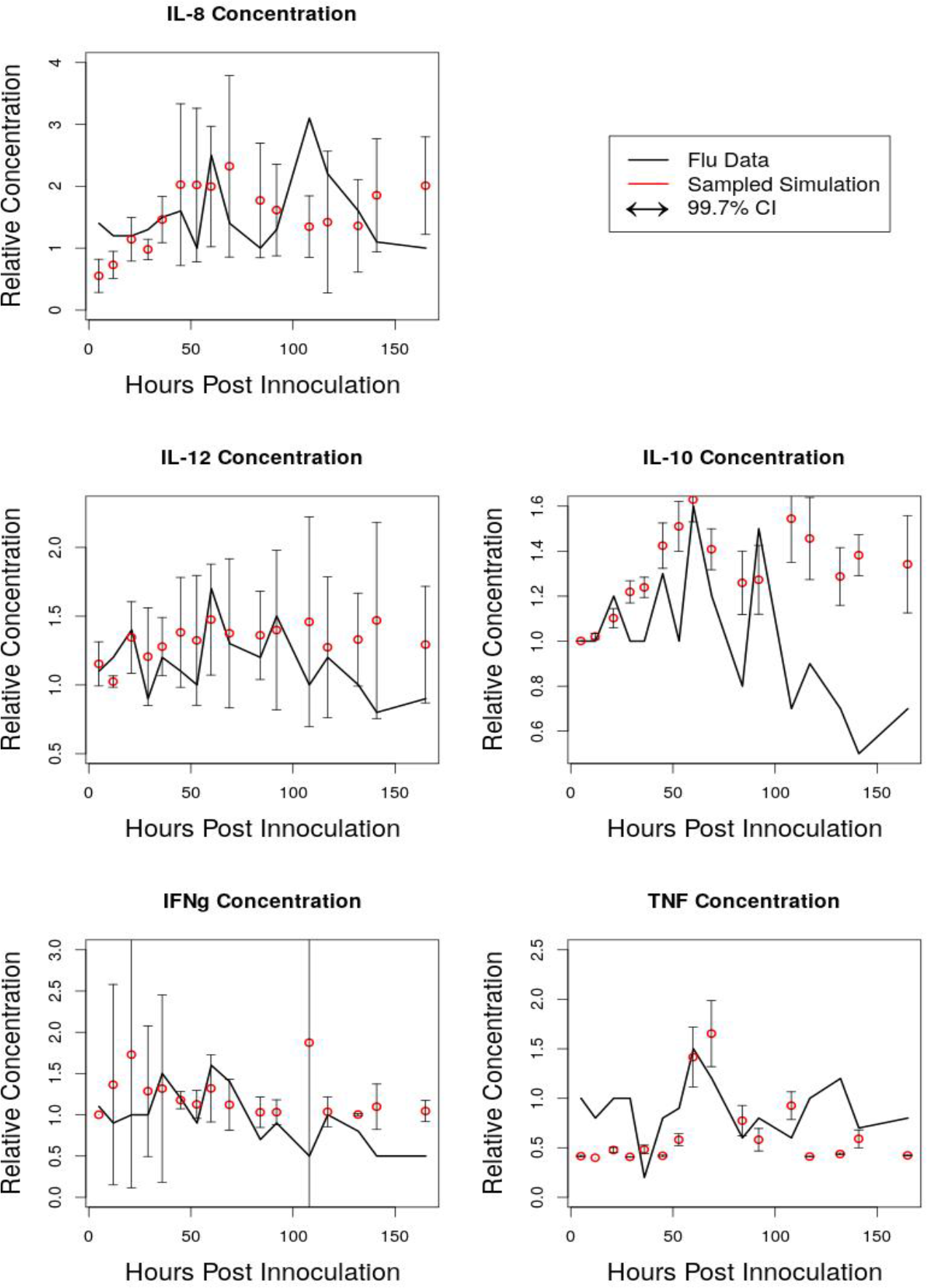
Plots of selected cytokine courses from flu challenge data compared to moderate infection courses

## 0.4.0 Discussion

The CIABM accurately models the general dynamics (cellular populations and cytokine time series) of an influenza infection. The CIABM generates dynamics which are similar to that of many different infections and display characteristics similar to those of the target real-world influenza challenge trial, displayed in Fig. 11. In Fig. 11 we compare the first 1400 steps of a simulated moderate infection to the cytokine data in McClain et al [21]. which tracks cytokines in a flu infection challenge. We can see that our range of cytokines presented within the model is similar to real world conditions. The dynamics shown in Figs. 3–10 show that in addition to cytokine courses being similar to real world measurements overarching immune dynamics are congruent with those seen in the real world [43–45]. While these dynamics are not perfect matches in every case the CIABM is extremely flexible as a modeling tool and can be further tuned.

The CIABM can also simulate asymptomatic infections, presenting similar, but blunted dynamics when comparing with more severe disease: the innate immune system responds first, the adaptive immune system takes several days to react, and the adaptive immune system develops long term immunity to the target pathogen. The flexibility of the CIABM to model a variety of infections in inherent in the design, and by expanding on previous work we believe we can quickly produce better calibrated dynamics. We have previously used machine learning on the meta-parameter matrix of the IIRABM to produce dynamics centered around observed data [10].

While the current paper presents simulations of influenza, the CIABM is designed as a general-purpose model of the human immune system able model a variety of infections. This can be done by adjusting the parameters associated with a particular virus, in terms of infectivity, replication rate and target cells. For instance, the current flu simulations do not allow the infection of immune cells; alternatively, viruses that do invade various immune cell types (such as Dengue or Ebola) can be given that property. Also, the current representation of the “tissue” layer is generically assigned to respiratory epithelium; these cellular agents, too, could be modified to represent other susceptible organs (such as intestinal or cardiac tissue). With a combination of existing mechanistic knowledge/hypothesized mechanisms regarding a virus and any data set regarding that infection the CIABM rule matrix can be tuned to model that infection. This can then be recalibrated with the addition of more data. For a single calibration target within the suite of targets we could use there are certainly many configurations of the meta-parameter matrix that would result in an acceptable match. By adding additional targets and additional data we can find a smaller set of matrices that fit all the targets simultaneously, and then use the calibrated model to understand the dynamics of that infection.

The CIABM is additionally flexible in that it is a framework to which other cell types and signaling and effector compounds can be added. The addition of natural killer T cells, plasmacytoid dendritic cells, B cells, and other cell types would be a straightforward process as the code is written in such a way that any cell type can be built from a set of generic functions, and any effector functions can then be incorporated through the meta-parameter matrix in a simple fashion. Epithelial cells additionally can be simply adjusted to produce differing compounds or function slightly different to represent different types of epithelial tissue.

We plan to adjust the CIABM to fit the known dynamics of SARS-CoV-19 infections in order to better understand viral dynamics. Because of the ability to use multiple calibration targets we can use many different data types to calibrate our model. By using a genetic algorithm, the meta-parameter matrix can be rapidly tuned to several calibration targets simultaneously, and then we can explore the factors that affect different disease outcomes. If some of these factors can be modulated with known medicines we may be able to inform new treatment regimens for SARS-CoV2-19 among other as-of-yet unknown diseases.

The rapid deployment and high degree of flexibility of the CIABM make it a highly useful tool for exploring emerging diseases without the need for expensive trials. By gathering simple diagnostic data, using single-point non-time-series calibration targets, and population scale statistics we can tune the model to any disease which primarily effects epithelial surfaces. By using the CIABM we can then explore viral dynamics, treatment regimes, and potential drug targets for viral diseases.

## Acknowledgements

This work was supported by the National Institutes of Health Grant #UO1EB025825.

