## Supplemental Material Table 1 for "The Cellular Immunity Agent Based Model (CIABM): Replicating the cellular immune response to viral respiratory infection"

| Rule Table for CIABM | | | | |
|---|---|---|---|---|
| **Cell Type** | **Rule in Model** | **Upregulation** | **Downregulation** | **Source of Base Rule** |
| Endothelial Cells | Activation | IL-1, IFNa | Lack of IL-1, IFNa | Introna & Mantovani (1997) |
| **The following rules only apply during endothelial activation** | | | | |
| Endothelial Cells | IL-8 production | Constant | Constant | Wolf et al (1998) |
| Endothelial Cells | PAF production | Constant | Constant | Bussolini & Camussi (1995) |
| Epithelial Cells | Activation | Viral particle intrusion | N/A | Mogensen & Paludan (2001) |
| Epithelial Cells | Viral Breakdown | Cytotoxins, IL-1 | N/A | Hsu & Spindler (2012) |
| Epithelial Cells | Viral Reproduction | Virions | Endotoxins and IFNa | Mahmoudabadi et al (2017) |
| Epithelial Cells | Endotoxin production | Cytotoxin, viral particles | Lack up upreg. | Lotz et al (2006) |
| Epithelial Cells | Tissue Damage leading to Cell Death | Cytotoxins, Endotoxins, Virions | N/A | Mehrbod et al (2019) |
| **The following rules only apply during epithelial activation** | | | | |
| Epithelial Cells | IFNa production | Constant | IL-10 | Mogensen & Paludan (2001) |
| Epithelial Cells | IL-1 production | Constant | Constant | Mogensen & Paludan (2001) |
| Epithelial Cells | TNF production | IL-2 | IL-10 | Mogensen & Paludan (2001) |
| Monocytes | Activation | PAF, TNF, endotox, IL-1 | IL-2 and IL-10 | Silva et al (2002) |
| Monocytes | IL-10 deactivation factor | TNF | N/A | An G. (2004) |
| Monocytes | IL-10 production | TNF, IL-1 | N/A | An G. (2004) |
| Monocytes | IL-1ra production | IL-1 | N/A | An G. (2004) |
| Monocytes | sIL1r production | IL-1r | N/A | An G. (2004) |
| Monocytes | sTNFr production | TNFr | N/A | An G. (2004) |
| **The following rules only apply during monocyte activation** | | | | |
| Monocytes | GCSF production | PAF, TNF, IFNg | IL-10 | An G. (2004) |
| Monocytes | IL-8 production | IL-1 | IL-2 and IL-10 | An G. (2004) |
| Monocytes | IL-12 production | TNF, IL-1, IL-8 | IL-10 | An G. (2004) |
| Monocytes | IL-1 production | PAF, TNF, endotox, IFNg, IL1r | IL-10 | An G. (2004) |
| Monocytes | TNF production | PAF, endotox, TNFr, IFNg | IL-2 and IL-10 | An G. (2004) |
| Monocytes | Virion removal, constant rate | Constant | Constant | Morgensen (1979) |
| PMN | IL1ra production | TNF, PAF | N/A | An G. (2004) |
| PMN | Programmed Cell Death | GCSF, TNF, IFNg | IL-10 | An G. (2004) |
| PMN | Migration into epi & oxidative burst | GCSF, TNF, IFNa, IL-1 | IL-2 and IL-10 | An G. (2004) |
| **The following rules only apply during an oxidative burst** | | | | |
| PMN | Cytotoxin production | TNF, IFNa, IL-1 | N/A | An G. (2004) |
| PMN | TNF production | Constant | Constant | An G. (2004) |
| PMN | IL1 production | Constant | Constant | An G. (2004) |

| Rule Table for CIABM continued | | | | |
| --- | --- | --- | --- | --- |
| **Cell Type** | **Rule in Model** | **Upregulation** | **Downregulation** | **Source of Base Rule** |
| Cytotoxic T | Activation | Antigen | N/A | Johnson et al (2003) |
| **The following rules only apply after Cyto T Activation** | | | | |
| Cytotoxic T | Cytotoxin production | Infected Cell Contact | N/A | Johnson et al (2003) |
| Cytotoxic T | Tissue Damage | Infected Cell Contact | N/A | Johnson et al (2003) |
| Myeloid DCs | Activation | Viral particles, GCSF | N/A | Cella et al (1997) |
| **The following rules only apply after mDC Activation** | | | | |
| Myeloid DCs | IL12 production | Constant | Constant | Gluckman et al (2002) |
| T Central Memory | Activation via antigen presentation by mDCs | mDC contact | N/A | Bouneaud et al (2005) |
| **The following rules only apply after mDC contact** | | | | |
| T Central Memory | IL2 production, constant upon activation | Constant | Constant | Sallusto et al (1999) |
| T Central Memory | Tem production | Constant | Constant | Sallusto et al (1999) |
| T Effector Memory | Activation | Viral particles | N/A | Sallusto et al (1999) |
| **The following rules only apply after Tem activation** | | | | |
| T Effector Memory | IFNg production | Constant | Constant | Sallusto et al (1999) |
| T Effector Memory | IL2 production | Constant | Constant | Sallusto et al (1999) |
| T Effector Memory | IL4 production | Constant | Constant | Sallusto et al (1999) |
| T Effector Memory | IL5 production | Il-2 | N/A | Sallusto et al (1999) |
| T Effector Memory | Proliferation into TEMRA | IL-2 | IL-5 | Dunne et al (2005) |
| T Effector Memory | TNF production | N/A | IL-10 | Sallusto et al (1999) |
| T Helper 0 | Development into TH1 or TH0 | Il-12, IFNg, IL-4 | N/A | Swain et al (2012) |
| T Helper 1 | IFNg production | TNF, IFNg, IL-1, IL-12 | N/A | Wagner et al (2010) |
| T Helper 2 | IL10 production | TNF, IFNg, IL-1 | Virions | Wang et al (2016) |
| T Helper 2 | IL4 production | N/A | IL-10 | Wang et al (2016) |
| TEMRA | Activation | Viral particles, GCSF | N/A | Henson et al (2012) |
| **The following rules only apply after TEMRA activation** | | | | |
| TEMRA | Cytotoxin production | N/A | IL-10 | Dunne et al (2005) |
| TEMRA | IFNg production | N/A | IL-10 | Dunne et al (2005) |

Table 1: Table of Rules for the CIABM. Contains activation rules, and subrules post-activation or during activation are denoted with a subsection surrounding them. In addition to any specific upregulation or downregulation factors, these rules are effected by the factors effecting activation.
